# MHC*IMP – Imputation of Alleles for Genes in the Major Histocompatibility Complex

**DOI:** 10.1101/2020.01.24.919191

**Authors:** David McG. Squire, Allan Motyer, Richard Ahn, Joanne Nititham, Zhi-Ming Huang, Jorge R. Oksenberg, John Foerster, Wilson Liao, Stephen Leslie

## Abstract

We report the development of MHC*IMP, a method for imputing non-classical HLA and other genes in the human Major Histocompatibility Complex (MHC). We created a reference panel for 25 genes in the MHC using allele calls from Whole Genome Sequencing data, combined with SNP data for the same individuals. We used this to construct an allele imputation model, MHC*IMP, for each gene. Cross-validation showed that MHC*IMP performs very well, with allele prediction accuracy 93% or greater for all but two of the genes, and greater than 95% for all but four.

## Introduction

The Major Histocompatibility Complex (MHC), located on chromosome 6 from 6p22.1 to 6p21.3, is the genetic locus most widely associated with human diseases (Price et al., 1999). This is likely due to the density of genes related to the immune system in the region. The MHC contains the Human Leukocyte Antigen (HLA) genes. HLA alleles are responsible for determining transplant compatibility, and have been found to be associated with numerous diseases and conditions, for example: autoimmune diseases (e.g. multiple sclerosis (Moutsianas et al., 2015; Sawcer et al., 2011), ankylosing spondylitis (Evans et al., 2011), psoriasis (Gudjonsson et al., 2003; Strange et al., 2010), rheumatoid arthritis (Han et al., 2014; Raychaudhuri et al., 2012)), communicable diseases (e.g. cerebral malaria (Hirayasu et al., 2012), HIV (Martin et al., 2007; Ramsuran et al., 2018)), cancer (e.g. Hodgkin lymphoma (Moutsianas et al., 2011), chronic lymphocytic leukemia (Gragert et al., 2014)), and adverse drug reactions (Bharadwaj et al., 2012).

Study of immunogenetic disease associations has been greatly facilitated by imputation methods for the alleles of the classical HLA genes: HLA-A, HLA-B, HLA-C, HLA-DPA1, HLA-DPB1, HLA-DQA1, HLA-DQB1, HLA-DRA, and HLA-DRB1. These methods allow alleles to be imputed on the basis of Single-Nucleotide Polymorphisms (SNPs). SNP identification using arrays remains significantly cheaper than Whole Genome Sequencing (WGS), and SNP-genotyped datasets thus continue to have larger sample sizes (Sudlow et al., 2015).

The first HLA imputation method, HLA*IMP (A. T. Dilthey et al., 2011; Leslie et al., 2008), has been widely used in association studies (Evans et al., 2011; Fairfax et al., 2012; Moutsianas et al., 2011; Sawcer et al., 2011; Strange et al., 2010). Other approaches have subsequently been developed, including HLA*IMP:02 (A. Dilthey et al., 2013), HIBAG (Zheng et al., 2013), SNP2HLA (Jia et al., 2013), and HLA*IMP:03 (Motyer et al., 2016).

To date, however, there are no methods for imputation of non-classical HLA and other MHC genes. These non-classical HLA and other genes are important. For example, HLA-E plays an important role in the recognition of cells by Natural Killer (NK) cells (Braud et al., 1998). HLA-E has been associated with diseases such as psoriasis (Zeng et al., 2013), bacterial infection (Tamouza et al., 2005), and leukemia (Xu et al., 2019). HLA-G expression has been associated with Crohn’s disease and ulcerative colitis (Rizzo et al., 2007). HLA-F has been associated hepatitis B and hepatocellular carcinoma (Zhang et al., 2012), and also with systemic lupus erythematosus (Jucaud et al., 2016). HLA-H variants are thought to be responsible for genetic haemochromatosis (Datz et al., 1997).

The MIC (Major Histocompatibility Class I Chain-Related) genes, of which MICA and MICB are the non-pseudogenes, are located in the MHC, but differ significantly from the classical HLA class I genes in their expression and products, and are highly polymorphic. They encode ligands for NK cell receptor NKG2D (Stephens, 2001). An association has been reported between MICA and MICB variants and enhanced susceptibility to leprosy (Tosh et al., 2006). They have also been associated with leukemia (Baek et al., 2018), and psoriasis (Choi et al., 2000; Romphruk et al., 2004).

The TAP (Transporter associated with Antigen Processing) genes TAP1 and TAP2, located in the MHC, have been associated with several diseases. For example, TAP1 and TAP2 polymorphisms have been associated with ankylosing spondylitis (Feng et al., 2009) An association with psoriasis has also been suggested (Pyo et al., 2003).

In this paper we present a method for imputing the alleles of multiple genes in the MHC. In particular, we focus on the imputation of non-classical HLA and other MHC genes (e.g. MICA, MICB, TAP1, and TA2). The classical HLA genes are also imputed. The method is based on prior work imputing classical HLA genes (A. Dilthey et al., 2013; A. T. Dilthey et al., 2011; Leslie et al., 2008; Motyer et al., 2016), and KIR gene copy number (Vukcevic et al., 2015). The imputation is done by using a random forest model (Breiman, 2001) to predict the allele for each gene for each haplotype.

## Reference Panel

### SNP Data

The SNP data used in the construction of the reference panel for this study was drawn from two sources, both reported in previous studies: “UCSF” (Yan et al., 2018) and “University of Dundee” (Nititham et al., 2018).

For all individuals from these sources, saliva samples were obtained. Both the psoriasis cases and healthy controls were genotyped on the Affymetrix UK Biobank platform. SNPs were called using Affymetrix Power Tools 1.18.0 using the parameters set out in the Affymetrix Best Practices Workflow document (Affymetrix, 2011)

This produced a panel of 3028 individuals in total, as shown Table 1.

**Table 1.** SNP data sources.

| Dataset | Number of Individuals |
| --- | --- |
| UCSF | 1523 |
| University of Dundee | 1505 |

### Allele calls

In order to build imputation models for the MHC genes, allele calls are needed for the individuals in the reference panel. 500 individuals were selected for sequencing, for most of whom we also had SNP data. These majority of these were individuals of European ancestry with a diagnosis of plaque psoriasis, as confirmed by a dermatologist, with a small number (15) of healthy controls.

The sequencing of the MHC region for these individuals was done using targeted sequencing by BGI Genomics. Alleles for genes in the MHC region were called from this sequence data using the SOAP-HLA program (Cao et al., 2013). These tools have previously been employed to develop an MHC reference panel of Han Chinese individuals (Zhou et al., 2016).

### Combining SNP data and allele calls

In order to create a reference panel for training the imputation models, we needed individuals for whom we had both SNP data and allele calls. There were 419 candidate individuals in the SNP datasets. After removing duplicates, triplicates, and first-degree relatives, 401 remained. These 401 individuals constitute the reference panel used to train the imputation models. A merged set of PLINK-format files was created with the unphased genotypes for these individuals (PLINK v1.90b6.9, (Purcell et al., 2007)).

### Phasing of alleles and SNP data

#### Allele encoding with PARSNPs

The imputation model requires phased data – we need to know which allele is associated with which haplotype (and thus particular SNP backgrounds). It is thus necessary not only to phase the SNP data, but to encode the allele information (as called from the sequence data) so that it is phased with the SNPs. This was done by encoding the alleles using a novel method: PARSNPs (Pseudo-Allele-Representing-SNPs). PARSNPs encode alleles using error-correcting codes, rather than the “one-hot” representation often used.

In a one-hot encoding, alleles are encoded using a string of bits which are all zero except for the *n*^th^ bit representing the presence of the *n*^th^ allele (a mapping from alleles to integers is required). This means that for all allele encodings that differ only in two bits – the vast majority of the bits are zero for all alleles. Phasing methods rely on switches that result in haplotypes not present in the data being phased (and possibly a reference panel) being improbable. One-hot encoding provides almost no such penalty: in most of the encoding of the allele, a switch from one long string of zeros to another is undetectable. This can result in the ones representing the two alleles for the individual being phased onto the same haplotype.

The motivation for PARSNPs is to minimise switch errors of this kind by using an allele encoding that guarantees that a switch in the middle of an encoded allele leads to haplotype that is not present in the data being phased – and thus recognizable as highly unlikely by the phasing algorithm. This is achieved by using error-correcting codes. Error-correcting codes allow a minimum Hamming distance (number of different bits) between valid codewords to be specified. As all allele encodings are valid codewords, a sequence of multiple switch errors would be required to lead to another bit sequence present in the data being phased. This reduces the chance of switch errors during phasing of the encoded allele.

The use of error correcting codes for allele encoding also allows phasing errors to be detected in multiple ways: switch errors often result in invalid codewords, as well as mismatches with known alleles for the individual.

Each field of the allele representation was encoded separately, using a 16-bit integer (two bytes) to represent each of the five possible allele fields (the last being alphabetic) in the current HLA nomenclature (Marsh et al., 2010). This representation is then encoded using an [8,4] Hamming code for each byte of each field, guaranteeing a Hamming distance of 4 between valid codewords (Hamming, 1950).^1^ This results in 32 bits per field, giving a 160 bit PARSNP representation of each allele.^2^ The binary representation can be mapped to DNA bases if that is required by the phasing software (e.g. 1 mapped to ‘A’, and 0 mapped to ‘G’).

The PARSNP representations of the alleles were embedded in the genotype data by finding the first location with a 160bp gap available (i.e. no SNPs in the data at those positions), starting from the centre of the gene. The PARSNPs are given SNP IDs of PARSNP*n* (where *n* is the bit number in [0, 160]) and SNP positions in bp in sequence from the insertion position. This approach guarantees no collisions between valid SNP IDs or SNP positions in the data, and that the PARSNP representation is embedded in the SNP background of the gene with which it should be phased.

#### Quality Control

Duplicate and monomorphic SNPs were removed before phasing, as were SNPs with more than 10% missing data.

#### Phasing

Phasing was then done with SHAPEIT v2.r904 (Delaneau et al., 2012). There is no external reference panel available for the non-classical HLA alleles used in this study. Given this, and the relatively small size of our dataset, it was expected that a significantly higher number of iterations than the default would be required for the phasing to converge. Varying numbers of iterations were tried, from the default of 20 through 80, 160, 320, to 3200. This was repeated 10 times for each number of iterations. Analysis showed that the rate of detectable phasing errors did not decrease beyond 320 iterations. Consequently, one of the 320 iteration phasing runs was picked at random for subsequent use.

### Creation of training data for imputation models

A separate training data set was constructed for each gene as follows. First, the allele for each haplotype was determined by decoding the embedded PARSNPs from the haplotype SNPs in the phased data. The PARSNPs were then removed from the SNP data. individuals with decoding errors missing allele calls, or alleles called at lower than two field resolution were removed from the training set for that gene.

The allele was then truncated to the desired resolution (two field) for imputation. Any genes found to be monomorphic at two field resolution were discarded. HLA-V was discarded as a result of this process.

For each gene, the SNPs in a window extending 50,000 bp either side of the start and end of the gene were extracted.^3^

This resulted in a training dataset with the properties shown in Table 2.

**Table 2.** MHC*IMP training data at two-field resolution.

| Gene | # of alleles | # of individuals | # of SNPs |
| --- | --- | --- | --- |
| HLA-A | 28 | 396 | 1590 |
| HLA-B | 49 | 392 | 2584 |
| HLA-C | 25 | 393 | 2690 |
| HLA-DMA | 2 | 396 | 1853 |
| HLA-DMB | 3 | 395 | 1944 |
| HLA-DOA | 2 | 397 | 1781 |
| HLA-DOB | 5 | 391 | 2261 |
| HLA-DPA1 | 6 | 397 | 1729 |
| HLA-DPB1 | 23 | 396 | 1731 |
| HLA-DQA1 | 16 | 396 | 2413 |
| HLA-DQB1 | 16 | 397 | 2433 |
| HLA-DRA | 2 | 397 | 2164 |
| HLA-DRB1 | 36 | 395 | 2355 |
| HLA-E | 2 | 397 | 1548 |
| HLA-F | 3 | 397 | 1568 |
| HLA-G | 6 | 383 | 1619 |
| HLA-H | 8 | 388 | 1626 |
| HLA-J | 2 | 397 | 1565 |
| HLA-K | 3 | 390 | 1615 |
| HLA-L | 2 | 397 | 1464 |
| HLA-P | 3 | 397 | 1614 |
| MICA | 10 | 174 | 2583 |
| MICB | 7 | 236 | 2489 |
| TAP1 | 5 | 397 | 2180 |
| TAP2 | 5 | 397 | 2239 |

#### Creation of cross-validation folds

In the absence of an independent reference dataset, *k*-fold cross-validation was used to evaluate the performance of our models (Devijver & Kittler, 1982). In *k*-fold cross validation, the dataset is split into *k* partitions (folds). Each fold is used as the test data for a model constructed using the other *k* − 1 folds as the training data.

To partition training data into *k* folds (we did three and five), for each MHC gene we first checked that we had at least *k* instances of each allele. If not, that gene would be excluded from the experiment. This is because we must not have monomorphic folds, and we would to have a chance of at least one instance of each allele in each fold. This bound guarantees that an assignment without monomorphic folds is possible, as is an allele instance in each fold (though of course this is not guaranteed). This criterion resulted in gene HLA-J being excluded at both the three- and five-fold levels, and gene HLA-DPA1 being excluded when five-fold cross-validation was used.

Individuals were then uniformly randomly assigned to one of the *k* folds. If any of the resulting folds was monomorphic, the assignment process was repeated until this was not so.

### Model Construction

For each gene, a random forest model (Breiman, 2001) was created to impute the allele for that gene for a haplotype. A random forest consists of a large number of decision trees (*ntree*), each of which is constructed using an independently drawn subset of training data (bagging). At each node of the tree, a number (*mtry*) of predictor variables (here SNPs) is considered in order to decide which branch to follow. Eventually a leaf of the tree is reached, which corresponds to the decision made (here the allele predicted). In order to make a prediction using a random forest, the predictions of all the trees in the ensemble are treated as votes for alleles. The fraction of votes for each allele can also be treated as a probability distribution.

We used the R *randomForests* package implementation of Breiman’s random forest model (Liaw & Wiener, 2002), with parameters:

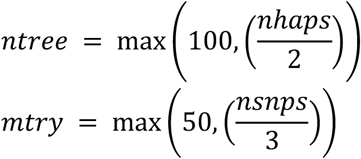

Where *nhaps* is the number of haplotypes, and *nsnps* the number of SNPs, in the training data for the gene. Experiments showed that performance was not sensitive to these parameters.

## Results and Discussion

We show and discuss results for five-fold cross-validation. In calculating accuracies for the model for each gene for each fold, “impossible” alleles (i.e. those not present in the training data for that fold) were excluded.

### Overall Performance

Figure 1 shows a summary of the overall allele prediction performance of our imputation models averaged across the *k* folds. The numerical data is shown in Table 3. Three performance measures are shown for the model for each gene: mean accuracy on the test data, mean accuracy on the training data, and average out-of-bag (OOB) accuracy, where accuracy is the percentage of alleles correctly predicted by the model from the SNP data. Genes are shown in order of increasing average number of alleles in the training data. We will focus on the accuracy on the test data in the discussion that follows. The performance on the training data is shown because it gives an indication of the extent to which the relationship between SNPs and alleles is “learnable” by the models.^4^ The average OOB accuracy gives an indication of the generalization performance of individual trees in the ensemble – the performance of the ensemble is expected generally to be better than that of individual trees.

**Figure 1.**
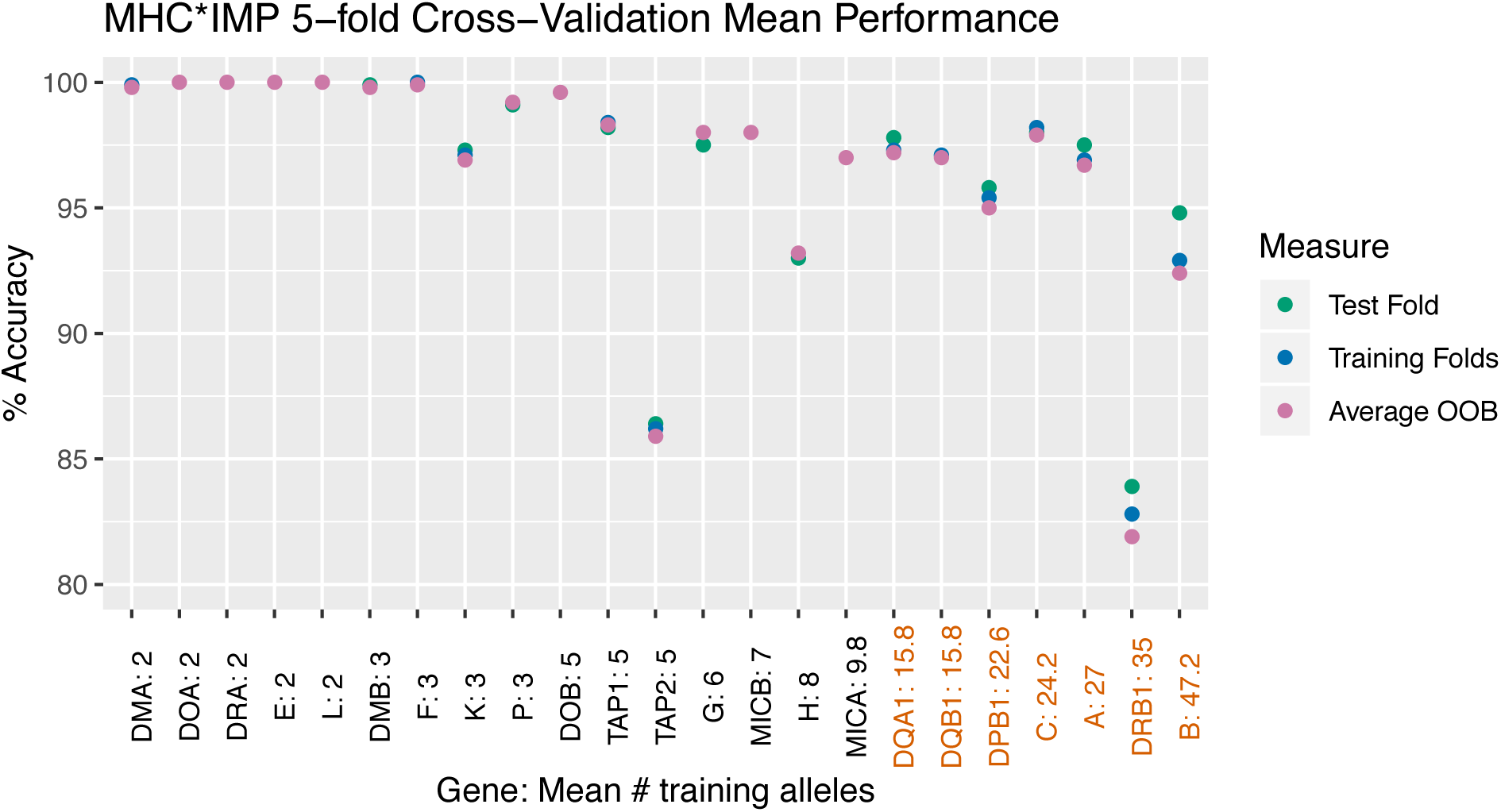
Overall Mean Cross-Validation Performance of MHC*IMP for each gene. Genes on the x-axis are in order of increasing average number of alleles in the training data for each of the five folds used for validation (that number is shown after the colon). The classical HLA genes, for which previous imputation methods exist, are shown in ochre; those from the extended MHC are in black. “Average OOB” indicates the average Out-Of-Bag error of individual trees in the random forest model for each gene.

Mean accuracy of MHC*IMP on the test data is 93% or greater for all but two of the 23 genes, and greater than 95% for all but four. In the context of previous methods for HLA imputation this is excellent performance given the size of the training set.

Performance generally decreases as the number of alleles increases, as would be expected – the larger than number of alleles, the fewer instances of each will appear in the training set. We would expect performance to improve when a larger reference dataset is available.

There are two outliers: HLA-DRB1 and TAP2, though even for these mean allele prediction accuracy is greater than 80%. HLA-DRB1 has been found to be one of the classical HLA genes more difficult to impute in previous studies (Motyer et al., 2016). It is possible that copy number variation in the HLA-DRB genes may confound either the allele calls from sequence data, or the SNP background in the neighbourhood of HLA-DRB1 (Doxiadis et al., 2012). TAP2 is discussed below.

Figure 2 shows the prediction accuracy of MHC*IMP on each of the five cross-validation folds, as well as the means that were shown in Figure 1. This gives an insight into the variance of the performance on the various folds. In general, the variance increases with the number of alleles. Again, HLA-DRB1 is an outlier, with by far the greatest variance. For all but the two outliers, HLA-DRB1 and TAP2, the accuracy on the worst fold is greater than 90%. In the application of MHC*IMP to a new data set, the entire reference panel would be used in training, and accuracy would be expected to be better than that obtained with *k* − 1 folds.

**Figure 2.**
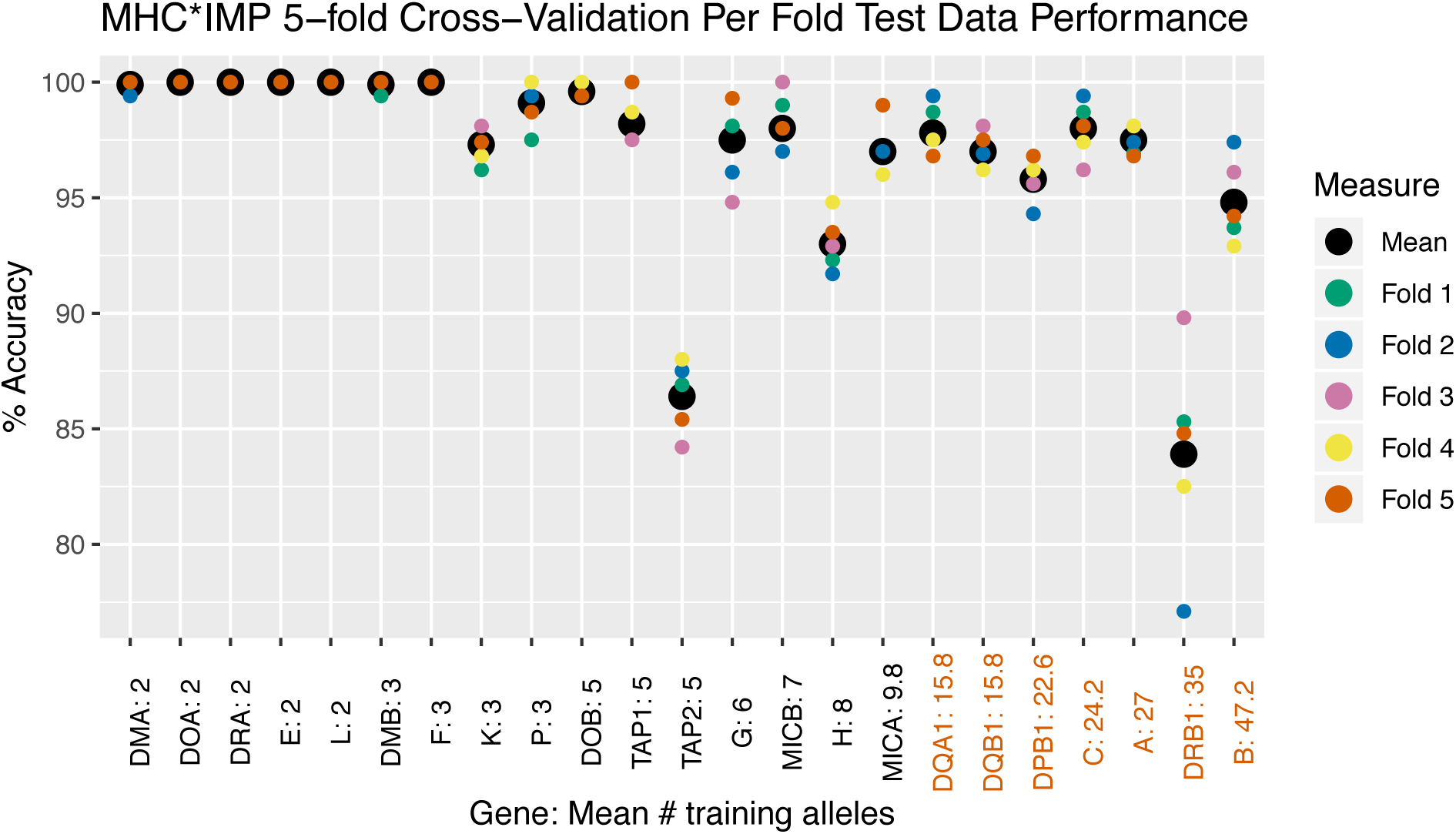
Performance of MHC*IMP on each of the five folds for each gene, and the means. Genes on the x-axis are in order of increasing average number of alleles in the training data data for each of the five folds used for validation (that number is shown after the colon). The classical HLA genes, for which previous imputation methods exist, are shown in ochre; those from the extended MHC are in black.

### Per Gene Performance

We will now discuss the performance on several individual genes in more detail. We will consider a “typical” case, HLA-C, and the two outliers, HLA-DRB1 and TAP2.

#### A typical case, HLA-C

Figure 3 shows the mean test and training set performance of MHC*IMP for gene HLA-C for each allele. Accuracy is excellent almost everywhere – the fraction of test instances correct (green) and training instances correct (blue) is close to 100%, except where there are very few instances of the alleles training data.

**Figure 3.**
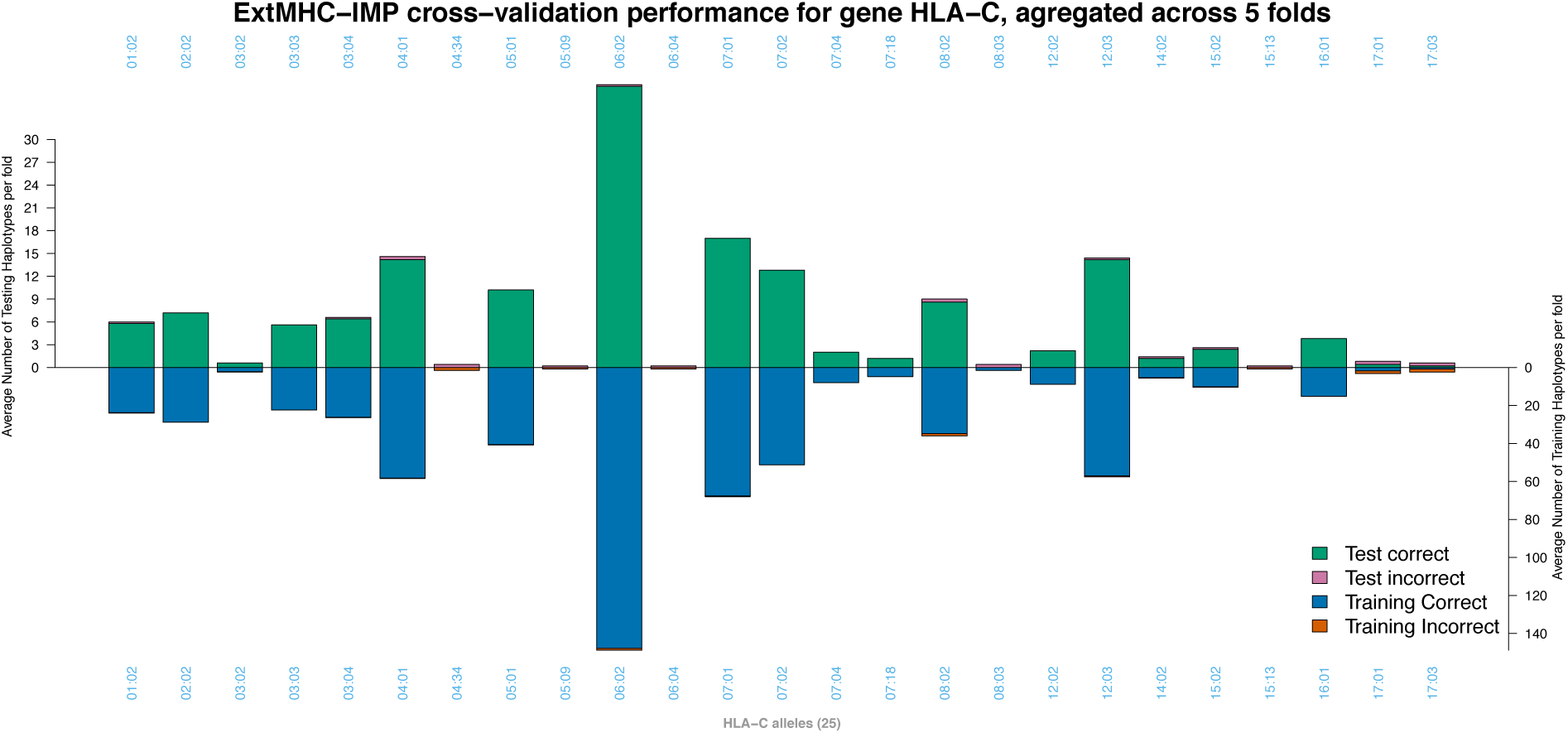
Mean per allele prediction performance for gene HLA-C.

Examining confusion matrices makes what is happening clearer. For example, Figure 4 shows the errors that were made for HLA-C when fold 4 was used as the test set:

**Figure 4.**
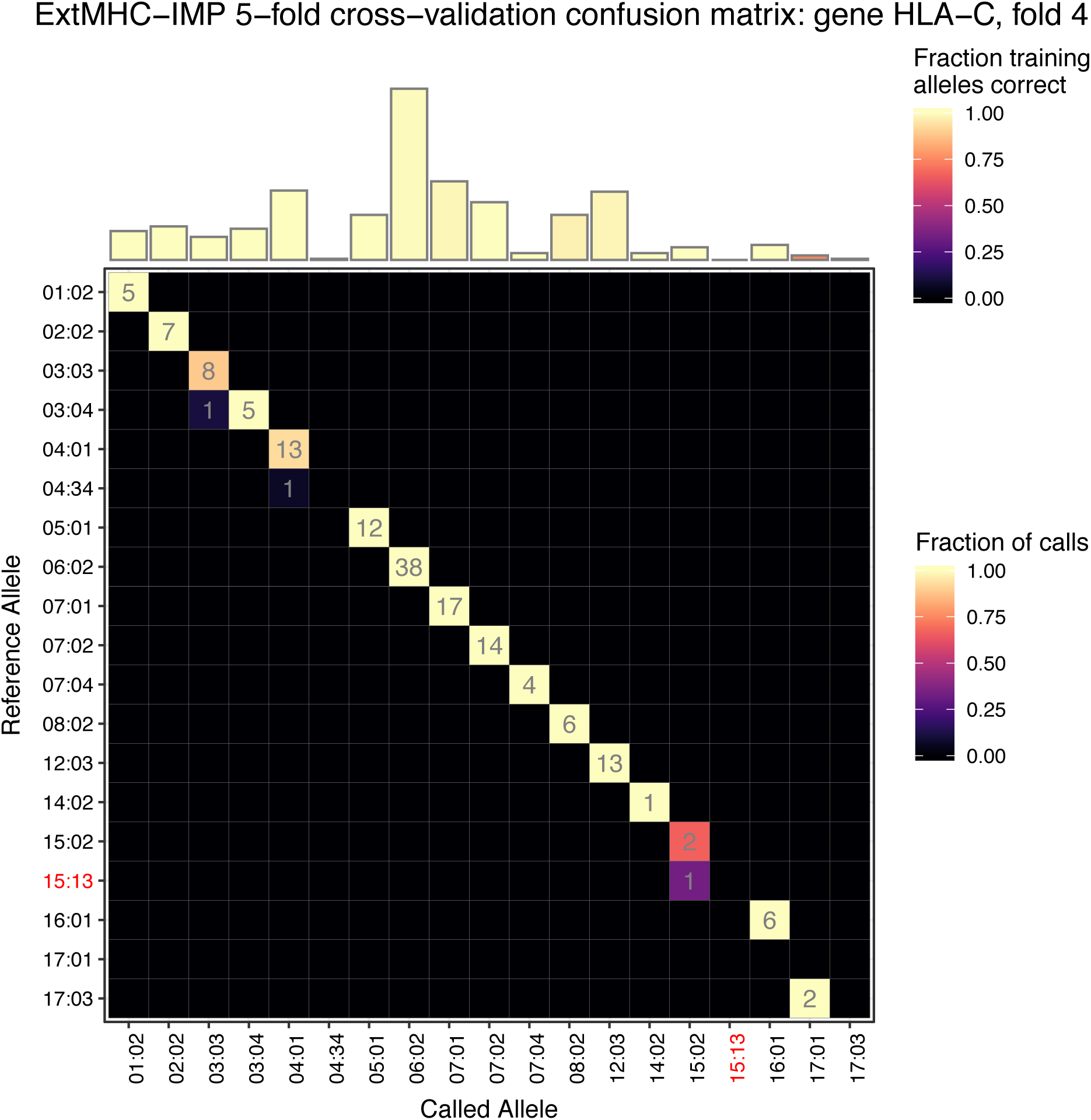
Confusion Matrix for gene HLA-C, with fold 4 as the test data. The histogram at the top of the figure shows the number of instances of each allele in the training data, the colour indicating the training set accuracy for that each allele. The matrix cells show the test data counts for each non-zero cell, with the colour indicating the fraction of called alleles corresponding to the known reference alleles for the test data. “Impossible” alleles are labelled in red (here allele HLA-C*15:13 was not present in the training data, and thus could not possibly be called correctly).

- One of the six HLA-C*03:04 instances is called as the more frequent HLA-C*03:03.
- The single HLA-C*04:34 instance is called as the relatively frequent HLA-C*04:01. There were extremely few instances of HLA-C*04:34 in the training data.
- The impossible HLA-C*15:13 instance is called as HLA-C*15:02.
- Both the HLA-C*17:03 instances are called as HLA-C*17:01. There were extremely few instances of either allele in the training data.

In summary, when the calls are wrong, they are still correct at one field resolution, and tend to the more common allele with that first field. Incorrect calls tend to occur when there are very few instances of the allele in the training data. Performance could thus be expected to improve with a larger reference panel.

#### A difficult case, HLA-DRB1

Figure 5 shows the mean test and training set performance of MHC*IMP for gene HLA-DRB1 for each allele. There are multiple alleles where performance is poor (i.e. a larger fraction of bar is pink (test data) or orange (training data). In all such cases, performance is (approximately) equally poor on both test and training data.

**Figure 5.**
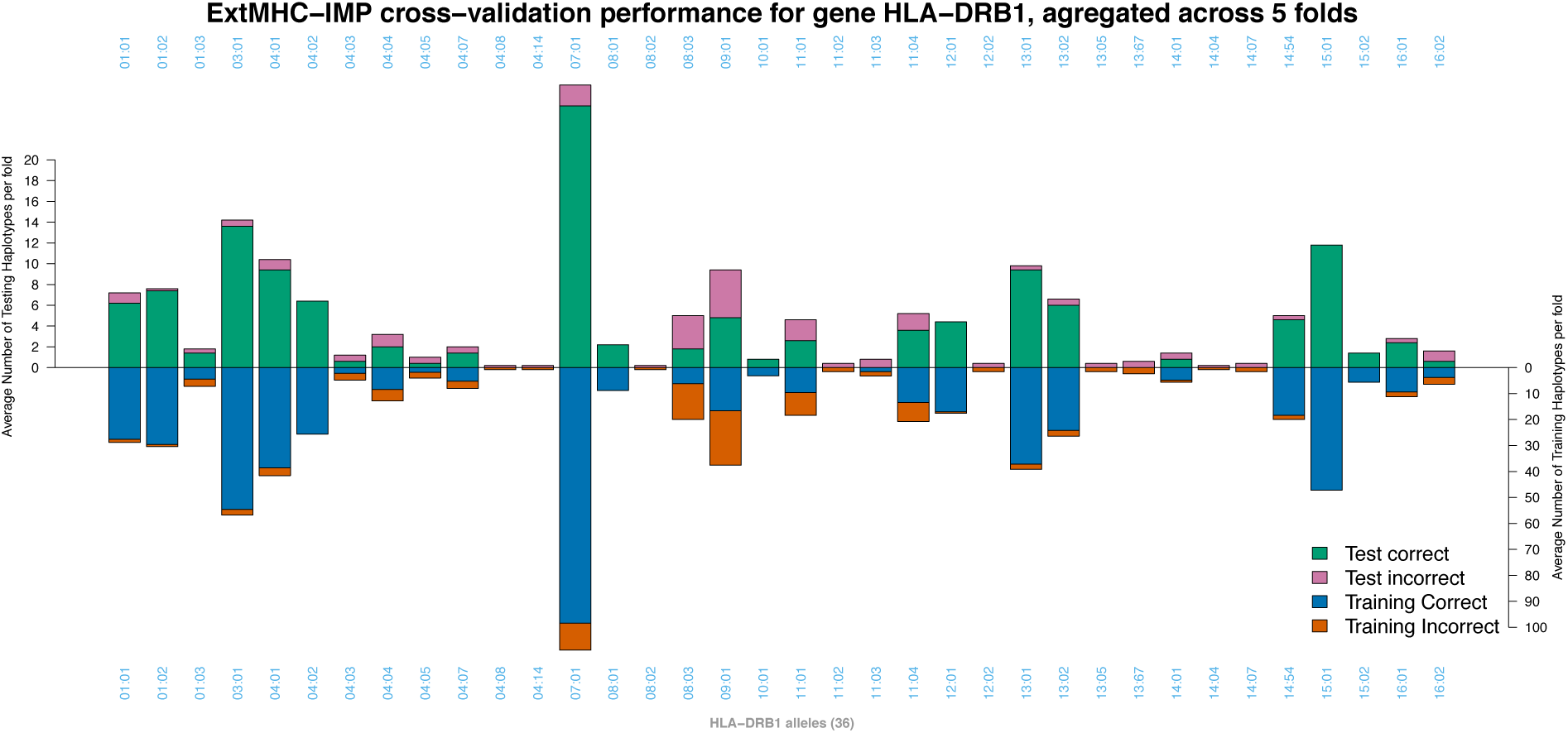
Mean per allele prediction performance for gene HLA-DRB1.

In this case, the confusion matrices shed little light – except that, as expected, poorer performance generally occurs for alleles with fewer training instances. Figure 6 shows the errors that were made for gene HLA-DRB1 when fold 2 was used as the test set. Whilst for some groups of alleles, incorrect calls are still correct at one field resolution (e.g. HLA-DRB1*14 and HLA-DRB1*16), this is not the case of the alleles HLA-DRB1*04, HLA-DRB1*07, HLA-DRB1*08, HLA-DRB1*09, HLA-DRB1*11, and HLA-DRB1*12.

**Figure 6.**
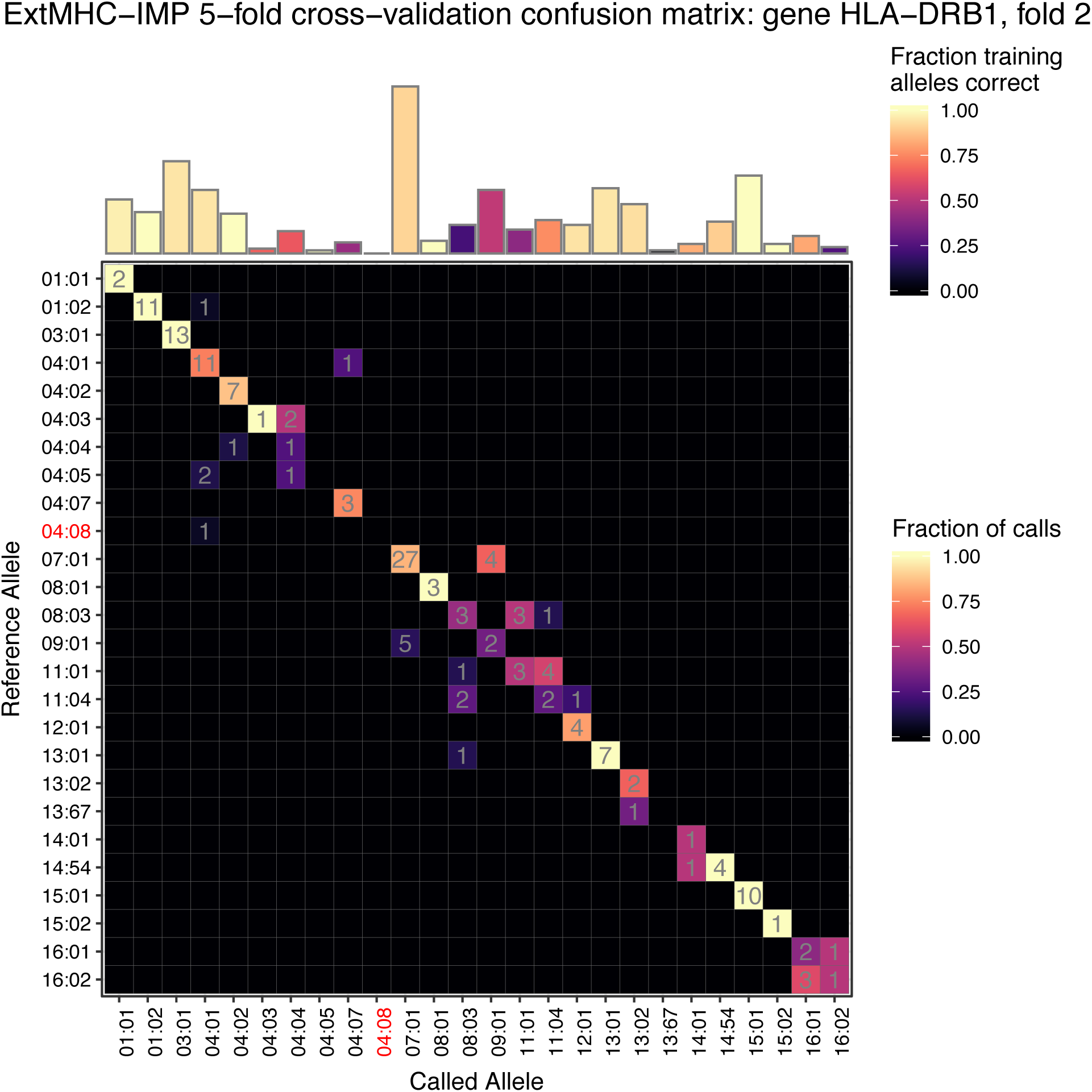
Confusion Matrix for gene HLA-DRB1, with fold 2 as the test data. See explanation of figure structure for Figure 4.

This suggests that either there is insufficient variation in the SNPs in reference panel for the model to capture the differences between these alleles, or that the calls for some alleles in the reference panel itself are incorrect.

#### A difficult case, TAP2

Figure 7 shows the mean test and training set performance of MHC*IMP for gene TAP2 for each allele. Performance on test and training data is again essentially identical. Allele TAP2*01:02 is evidently very hard to learn from our reference panel. Allele TAP2*01:03 is rare and apparently unlearnable here.

**Figure 7.**
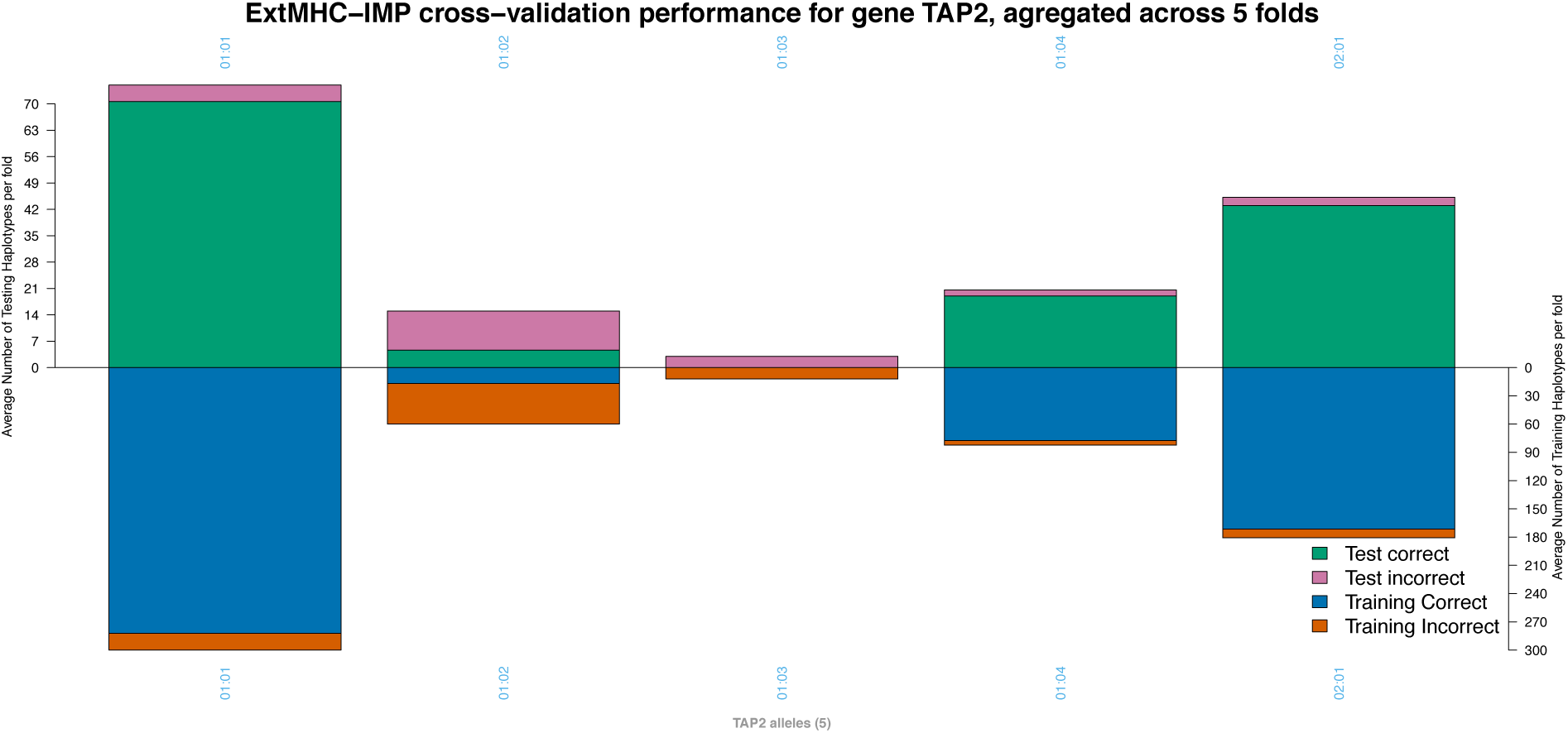
Mean per allele prediction performance for gene TAP2.

Figure 8 shows the errors that were made for gene TAP2 when fold 2 was used as the test set.

**Figure 8.**
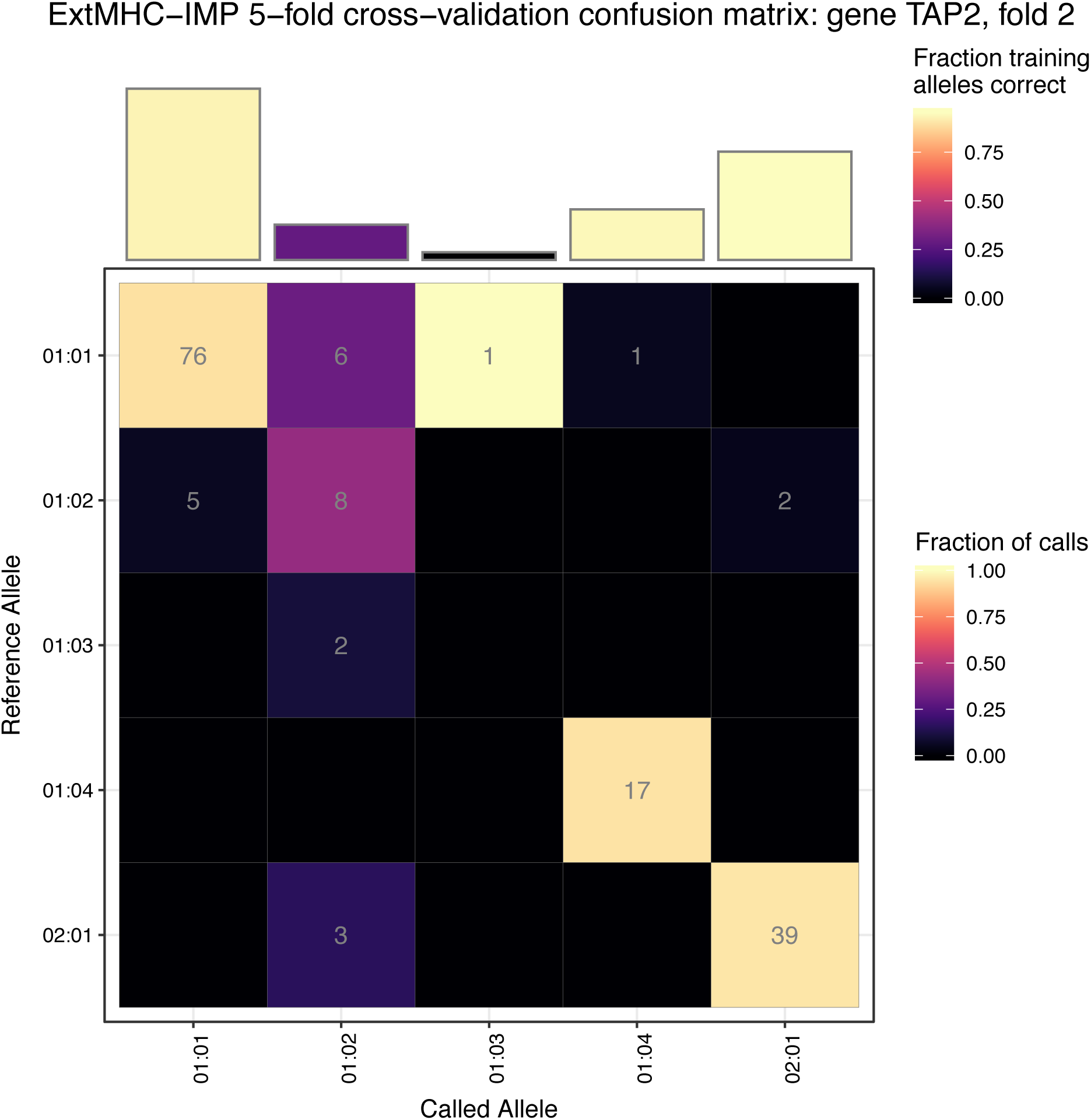
Confusion Matrix for gene TAP2, with fold 2 as the test data. See explanation of figure structure for Figure 4.

- 5 of the 13 test instances are called as the much more common TAP2*01:01
- the 2 TAP2:01:03 instances are called as TAP2:01:02, as are 3 of the 42 TAP2*02:01 instances
- 2 TAP2*01:02 instances are called as TAP2*02:01 – the confusion goes in both directions

It is difficult to draw conclusions here, except that the great majority of instances of the common alleles (TAP2*01:01, TAP2*01:04, and TAP2*02:01) are called correctly. TAP*01:02 is decidedly odd: it is the most common allele for others to be called as incorrectly, and yet it is only called correctly itself ∼50% of the time. We can only conclude that our reference panel contains insufficient, or even contradictory, information for this gene.

## Conclusion

We created a reference panel consisting of 401 individuals for 25 genes in the MHC using allele calls from WGS data, combined with SNP data for the same individuals. We used this to construct an allele imputation model, MHC*IMP, for each gene. Cross-validation showed that MHC*IMP performs very well, with allele prediction accuracy 93% or greater for all but two of the genes, and greater than 95% for all but four. We expect the performance of the MHC*IMP approach to improve still further when a larger reference panel is available.

## Acknowledgements

This work was supported in part by funding from the NIH to W.L. (5U01AI119125).

## Supplementary Data

**Table 3.** Overall Mean Performance of MHC*IMP Models.

| Gene | Training data accuracy | Test data accuracy | Average OOB accuracy | # Training alleles | # Testing alleles |
| --- | --- | --- | --- | --- | --- |
| HLA-A | 96.9 | 97.5 | 96.7 | 27 | 20 |
| HLA-B | 92.9 | 94.8 | 92.4 | 47.2 | 33.6 |
| HLA-C | 98.2 | 98 | 97.9 | 24.2 | 18.4 |
| HLA-DMA | 99.9 | 99.9 | 99.8 | 2 | 2 |
| HLA-DMB | 99.8 | 99.9 | 99.8 | 3 | 3 |
| HLA-DOA | 100 | 100 | 100 | 2 | 2 |
| HLA-DOB | 99.6 | 99.6 | 99.6 | 5 | 5 |
| HLA-DPB1 | 95.4 | 95.8 | 95 | 22.6 | 18.8 |
| HLA-DQA1 | 97.3 | 97.8 | 97.2 | 15.8 | 13.6 |
| HLA-DQB1 | 97.1 | 97 | 97 | 15.8 | 14.8 |
| HLA-DRA | 100 | 100 | 100 | 2 | 2 |
| HLA-DRB1 | 82.8 | 83.9 | 81.9 | 35 | 26.8 |
| HLA-E | 100 | 100 | 100 | 2 | 2 |
| HLA-F | 100 | 100 | 99.9 | 3 | 2.8 |
| HLA-G | 98 | 97.5 | 98 | 6 | 5.6 |
| HLA-H | 93.2 | 93 | 93.2 | 8 | 7.4 |
| HLA-K | 97.1 | 97.3 | 96.9 | 3 | 3 |
| HLA-L | 100 | 100 | 100 | 2 | 2 |
| HLA-P | 99.2 | 99.1 | 99.2 | 3 | 3 |
| MICA | 97 | 97 | 97 | 9.8 | 8.8 |
| MICB | 98 | 98 | 98 | 7 | 6.4 |
| TAP1 | 98.4 | 98.2 | 98.3 | 5 | 4.6 |
| TAP2 | 86.2 | 86.4 | 85.9 | 5 | 5 |

## Footnotes

1 Encoding each field separately is simple and easy to interpret, but has the disadvantage that switch errors at field boundaries are not greatly penalized – a switch to another valid codeword (encoding a different value for the following field) is possible. Indeed the few errors observed in allele phasing were much more frequently of this type than a mid-codeword switch, as expected. This issue could be address by using an encoding with a single codeword for each allele, with a specified minimum Hamming distance between valid codewords. There would thus be no positions at which a switch would not be penalised. Reed-Solomon Codes provide one way of doing this (Reed & Solomon, 1960).

2 This is more compact than a one-hot representation if there are more than 160 possible alleles – which is the case for some HLA genes. Moreover, the representation is unique – it does not depend on how many alleles are present in the particular dataset (as one-hot encoding typically does). If there are no alleles with the numeric value of a field greater than 255, the most significant byte will be constant, and thus removed before phasing as monomorphic.

3 Larger window sizes were investigated, but the same – or sometimes worse, for very large windows – performance was observed. This is perhaps unsurprising with a small training set and a complex model capable of learning spurious correlations between alleles and distant SNPs.

4 The fact that test fold performance is most frequently better than training set performance may be an artefact of the fact that rare (and thus hard to learn) alleles are more likely be present in the larger training dataset (the number of haplotypes is perhaps small enough that the fact that there can’t be fractional instances matters). If all instances of a rare allele end up in training set, they are excluded from test set performance calculations.

